# Differential contributions of nuclear lamina association and genome compartmentalization to gene regulation

**DOI:** 10.1101/2022.09.12.507606

**Authors:** Priyojit Das, Rebeca San Martin, Rachel Patton McCord

## Abstract

Interactions of chromatin with the nuclear lamina play a significant role in properly organizing the genome in 3D space and in regulating gene expression. Genome wide studies have inferred the global association between the lamina, heterochromatin, gene repression and the B genomic compartment, and repositioning genes to the lamina can result in their repression. However, there are scenarios in which these features are discordant and, in those cases, the relative contribution to gene regulation of genomic compartment, chromatin, and lamin association status can be examined. Here we compared datasets from cell lines representing different states of differentiation across different cell type lineages to examine the relationships between changes in genomic compartmentalization, lamin association, and gene expression. With these data, we could examine, for example, what gene expression changes occur when a B compartment region is moved from the nuclear interior to the nuclear lamina and what differences exist between lamin associated and internal A compartment regions. In general, we observed an additive rather than redundant effect in which lamin association and compartment status both contribute to gene expression state. However, we found that cell type lineages differed in whether compartment status or lamin association had a dominant influence on gene expression. Finally, we identified conserved trends of how compartment and lamin association status influence the likelihood that gene expression will be induced or repressed in response to a physiochemical treatment.

## INTRODUCTION

Inside the nucleus, chromosomes interact among themselves, with other different nuclear substructures and the nuclear lamina, leading to the formation of 3D spatial genome organization^1^. Several aspects of this organization have been implicated in gene regulation^2–6^, including a repressive role of both chromatin tethering to the nuclear lamina and spatial segregation into the B compartment.^5,6^ The nuclear lamina is a protein meshwork underneath the nuclear membrane which contributes to the structural stability of the nucleus^7, 8^. The chromatin regions tethered to the lamina (lamina-associated domains; LADs) are typically between 0.1 and 10 Mb in size and are generally gene-poor, transcriptionally repressed, heterochromatin-rich regions^6^. Several studies have shown that tethering an active region to the lamina tends to lead to decreased expression of the genes located in that region^9–13^. Targeted activation or inactivation of genes can promote chromosome region detachment from or attachment to the nuclear lamina, respectively.^14^ However, it is not clear what purpose would be served by relocating an already inactive region to the lamina or an already active region toward the interior. Genome-wide chromosome conformation capture experiments (Hi-C) have demonstrated that chromosomal regions spatially segregate into different compartments in the nucleus^15, 16^. The regions classified as belonging to the B compartment are enriched in heterochromatic marks, gene-poor regions, and inactive genes and are often associated with the nuclear lamina. Regions that switch into the B compartment often experience gene repression and vice versa^17^. But, despite the frequently reported correlations between compartment status, lamin association, and gene expression, the combinatorial effects of different repressive or activating environments on gene expression are not fully understood. How are genes affected when a typically active A compartment region is located at the lamina? Does an already repressed B compartment region become further stably repressed when located at the nuclear periphery as compared to the interior? Detailed analyses of the relationships between lamina association, gene expression, and compartmentalization are often performed in a single cell type, but does the relative contribution of these factors vary by cell type or cell lineage?

One approach to shed light on these questions is to consider cells across the spectrum of differentiation status in different cell type lineages. As a stem-like cell progresses through differentiation, finally committing to a particular fate, gene expression, genomic compartmentalization, and LAD organization also change^18–24^. Thus, in this work, we integrated publicly available genomic data to compare cell lines in different lineages and stages of differentiation to investigate the relationship between compartment identity, LAD reorganization, and gene expression differences between cell lines.

Genome structure and lamina association can influence not only the stable expression or repression of genes in a particular cell type but also how likely a gene is to be induced by a stimulus^25, 26^. Given the same perturbation, different cell types will activate or repress different genes^27^, and we hypothesize that the pre-existing compartment and lamin organization state of the chromatin contributes to this differential gene expression. Thus, we also examined the impact of compartment status and lamina association on gene expression changes after stimulus or perturbation.

In this study, we present evidence of the existence of both conserved and lineage-specific trends of gene expression regulation. We found that there is most frequently an additive effect of LAD and compartment status on gene expression. However, for specific regions, depending on their underlying compartment identity and lamina association in different cell type lineages and states, we observed differences in whether compartment or LAD status dominate the gene expression effect. Further, across different cell types and perturbations, genes that were either induced or repressed were tended to be enriched in the interior A compartment state, suggesting that this positioning is most favorable for altering gene expression in either direction.

## RESULTS

### Chromosome compartmentalization and lamina association segregate cells by lineage

To investigate how the LAD reorganization contributes to the regulation of gene expression located in the different compartments of the genome, we required data representing: a) genome-wide lamina association (to assess whether a region is associated with the lamina), b) compartmental organization (to assess whether a region belongs to A or B compartment) and c) transcriptomics (for gene expression) data. Therefore, we selected cell types for which publicly available lamina association (mainly DamID data, unless otherwise mentioned), genome organization (Hi-C) and gene expression (RNA-seq) data were available. In total, eleven different cell types consisting of both primary cells and cell lines were collected for this study and classified based on developmental lineage and malignancy status based on known characteristics. Considering embryonic stem cells (H1ESC) as the node of origin, cells were separated into three lineages: a) mesoderm, derived from mesenchymal stem cells (HMSC), b) endoderm, primarily composed of epithelial cells and c) hematopoietic cells (Fig 1a) ^28, 29^. The mesoderm-derived cell lines were sub-divided into two families: fibroblasts (WI38, IMR90, Tig3, and Hffc6) and osteoblast-derived osteosarcoma (U2OS)^30–33^. Among the fibroblast cells, WI38, IMR90, and Tig3 are all female fetal lung fibroblast cell lines and, for the purpose of this analysis, considered interchangeable ^31–33^. The epithelial cell lines were classified as either potentially secretory (colon carcinoma – HCT116) or non-secretory (RPE)^34–36^. A special case of epithelial fibrosarcoma (HT1080) was considered a potential midpoint between a bona fide sarcoma (U2OS) and an epithelial line^37^. Finally, the hematopoietic cell lines were classified as either derived from a myeloid progenitor (K562 and HAP1) or a lymphoid progenitor (Jurkat)^38–40^.

**Figure 1.**
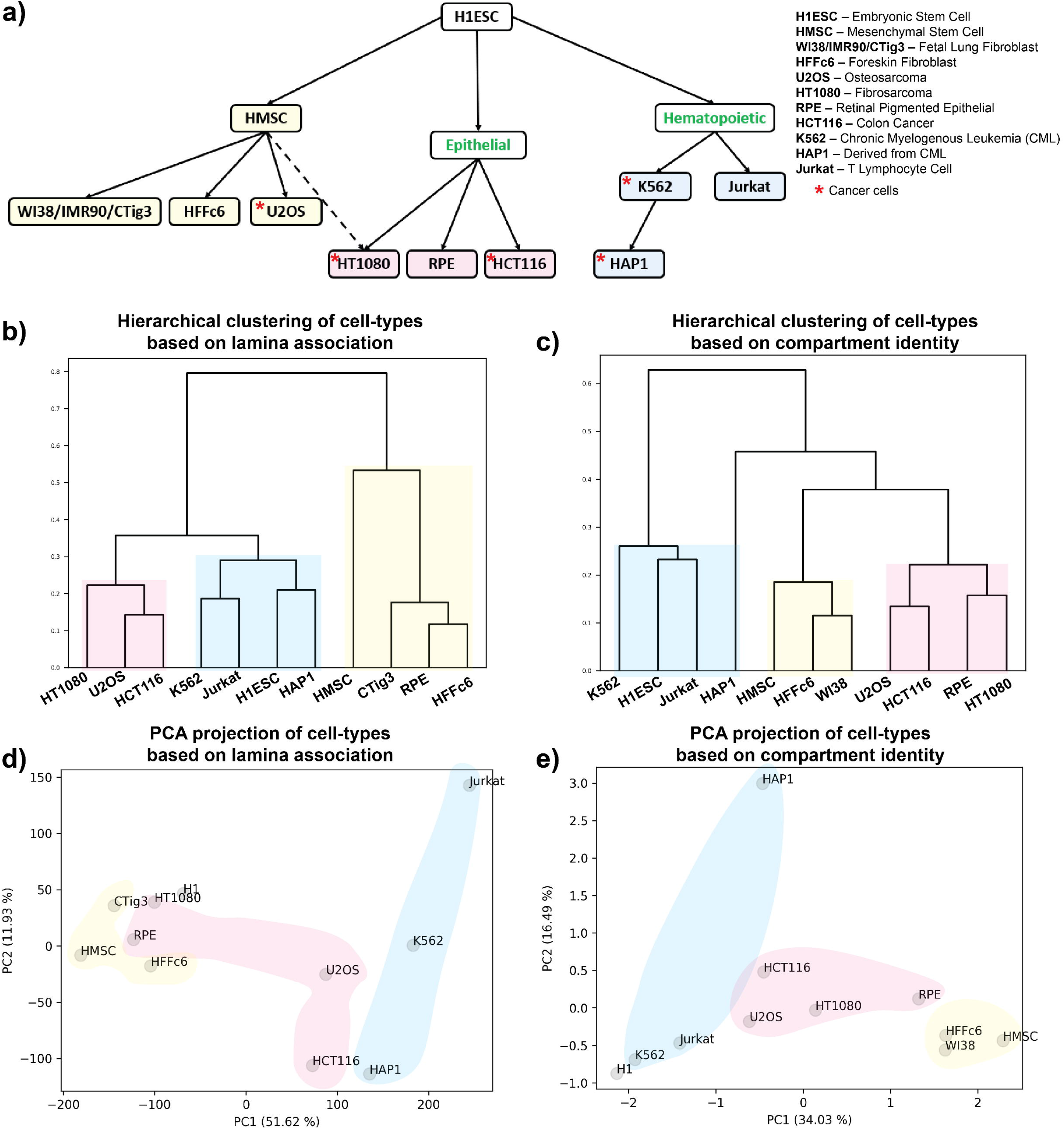
Cell lineage relationships are detected in both compartmentalization and lamin association patterns. **a)** Human primary cells and cell lines are classified based on their developmental lineage relationships and malignancy status based on known characteristics. Yellow: mesenchymal, pink: epithelial, blue: hematopoietic, green labels indicate intermediate states in lineages which do not have specific samples representing them. **b, c)** Hierarchical clustering of genome-wide lamina association (b) and compartment identity (c) data for different cell types at 40 kb and 250 kb resolutions respectively. Sub-trees are colored based on their lineages as shown in (a). **d, e)** Principal component analysis of genome-wide lamina association (d) and compartment identity (e) data for different cell types at 40 kb and 250 kb resolutions respectively. Cells are colored based on their lineages as shown in (a).

Once we had ordered the cell types based on prior domain knowledge, we compared this ordering to the unsupervised classification of the cell types based on either the lamina association or compartment identity data as described in the **Methods**. Briefly, for lamina association data, each 40 kb genomic region was assigned a lamina association value, where a value greater than zero signifies lamina association and vice versa. For compartmental organization, positive and negative values assigned to each 250 kb region indicate A and B compartment status respectively. In the case of the lamina association, hierarchical clustering produced a cell type grouping similar to what we anticipated based on published cell characteristics, with a few interesting exceptions (Fig. 1b). We found that U2OS, which is considered epithelioid-like osteosarcoma, presents a similar type of lamina association to bona fide epithelial cells. Surprisingly, the RPE (retinal pigment epithelial) cells grouped with the mesenchymal lineage cells rather than with other epithelial cells. This may be explained by the relatively small nuclear size and the incidence of polynucleated cells in RPE^41, 42^. Compartment organization clustering produced similar cell type ordering trends (Fig. 1c). In both datatypes, we observed H1ESC clustering with the hematopoietic lineage cells (Fig. 1b and 1c). Interestingly, principal component analysis (PCA) on both types of genomic data, showed that the principal component 1 (PC1) axis ordered the cell types based on their lineages, where specifically the mesenchymal and hematopoietic lineages can be found at the ends of the spectrum, with cells of epithelial lineage in the middle (Fig. 1d and e). This recapitulates the relationships observed by unsupervised clustering and suggests that mesenchymal and hematopoietic lineages differ significantly in both lamina and compartmental organization. Furthermore, we observed that cell types segregated according to the overall proportion of their genome associated with the lamina or B compartment (SFig. 1). Notably, H1ESC cells show a similar proportion of genomic regions with peripheral localization as the hematopoietic lineage cells. Similarly, U2OS and RPE cells show a similar global level of lamina association to epithelial and mesenchymal cells respectively (SFig. 1a). Overall, we find that lamina association and genomic compartmentalization vary in a lineage-specific manner.

### Lamina association adds another layer of gene regulation to compartment identity

We next evaluated how gene expression changes as active or inactive regions alter their spatial position inside the nucleus. We compared pairs of cell types that share a directional connection in our cell lineage tree (Fig. 1a). For the analyses, we started by comparing regions that keep their compartment identity unaltered but change their lamina association among the compared cell types. We then checked the gene expression fold change between cell types within those regions (Fig. 2 and SFig. 2). We found that genomic regions that have a similar compartment identity between cell types exhibit decreased or increased expression depending on their lamina association. Further, the association between lamin state and gene expression follows more strongly in the direction that would typically be associated with the underlying compartment state: gene upregulation is more likely to accompany a loss of lamina association for regions already in the A compartment and, conversely, gene repression is more likely to accompany relocation to the nuclear lamina for regions already in the B compartment. In contrast, a region that stays in the B compartment but shifts away from the lamina does not necessarily also show gene activation. Overall, these results suggest that the lamin association status of a genomic region further modulates the gene expression profile of that region, though the compartment identity remains unaltered.

**Figure 2.**
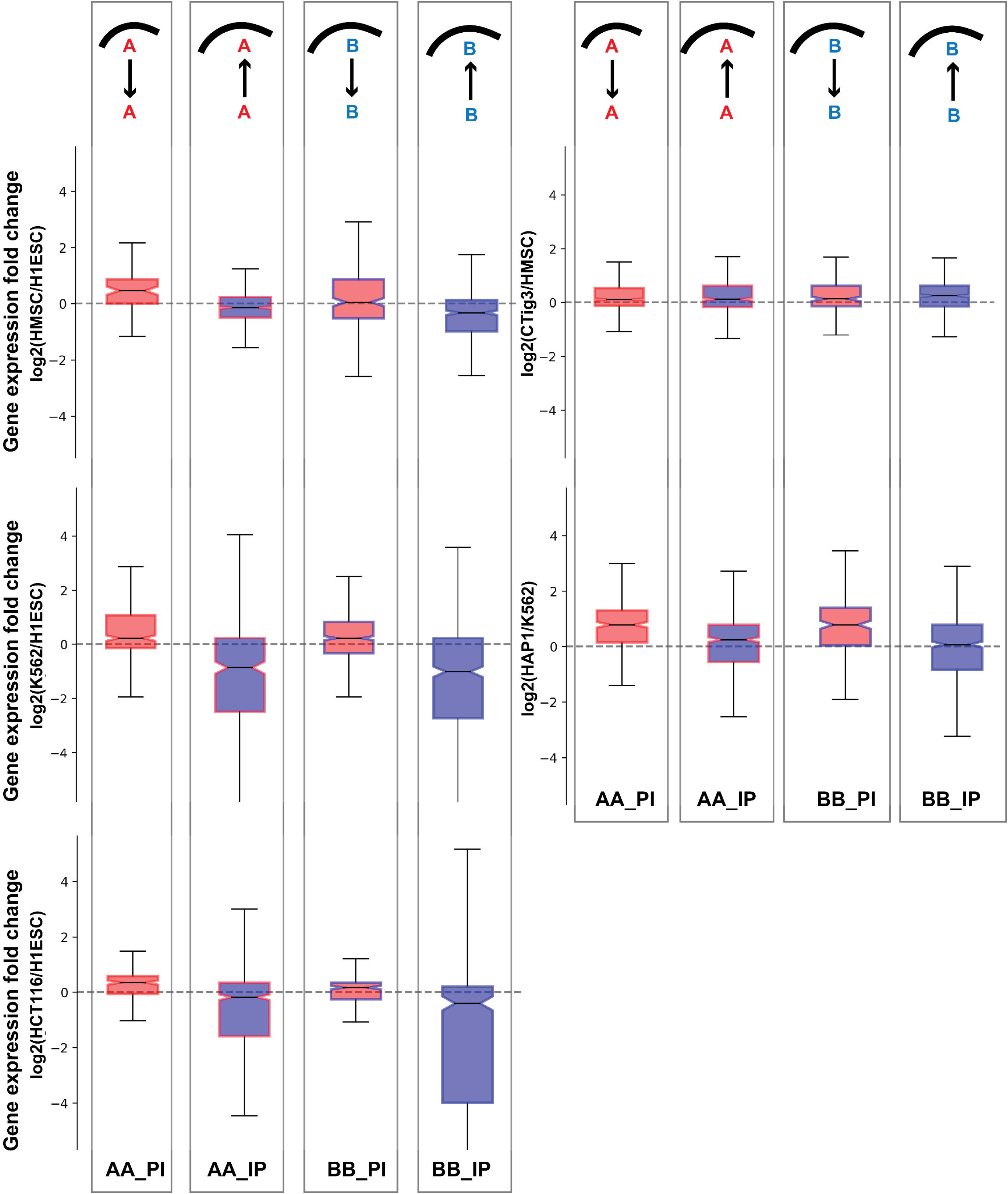
Gene expression change between cell types follows alterations in lamina association for genomic regions with unchanged compartment identity. Cell types being compared are indicated in y-axis labels. Comparisons are represented using acronyms. For example, ‘AA_IP’ indicates regions which belong to A compartment in both cell types but switch from no lamina association (“Internal”) to lamin associated (“Peripheral”) from the first to the second cell type. The compartment identity of each set of regions is also shown by box plot border color (red = A, blue = B) while the internal shading indicates the direction of lamin association switch (red = shift to non-LAD, blue = shift to LAD). The transition from one cell type to another for the set of regions is also shown in a cartoon over the boxplot where the curved line indicates the nuclear lamina. Boxplots show minimum, 1^st^ quartile, median, 3^rd^ quartile, and maximum.

### Coupling and decoupling of lamina association and compartmentalization effects on gene expression

We next extended our gene expression analysis to all possible combinations of lamina association and compartment identity alterations for each of the cell type comparisons (Fig. 3 and SFig. 3). For convenience, we refer to comparisons by acronyms such as BA_PI, indicating the compartment identity in each cell type (BA = B in the first cell type and A in the second cell type) as well as the LAD status in each cell type (PI = peripheral, LAD associated in the first cell type and internal, non-LAD associated in the second cell type). These comparisons further confirm that when compartment identity remains unaltered, changes in gene expression follow the lamina association. Since B compartment regions are often lamin associated and vice versa, we refer to B/P and A/I associations as “congruent”. In all cases where the second cell type in the comparison showed congruent compartment and LAD status, (e.g., AA_PI, BA_II, AB_PP, and BB_IP), gene expression changes follow LAD status changes. On the contrary, in regions for which lamina association is incongruent with compartment identity (e.g., AB_II, BA_PP, AA_IP, BB_PI, AA_PP, AB_PI, BA_IP, and BB_II) some cell types appear to be more governed by LAD type while others favor compartment identity control. For example, in the case of mesenchymal lineage cells and RPE cells, the gene expression of these incongruent regions matches the compartment identity more than the lamina association (SFig. 3). In contrast, the incongruent regions in comparisons among all other lineages show gene expression changes matching the lamin association status.

**Figure 3.**
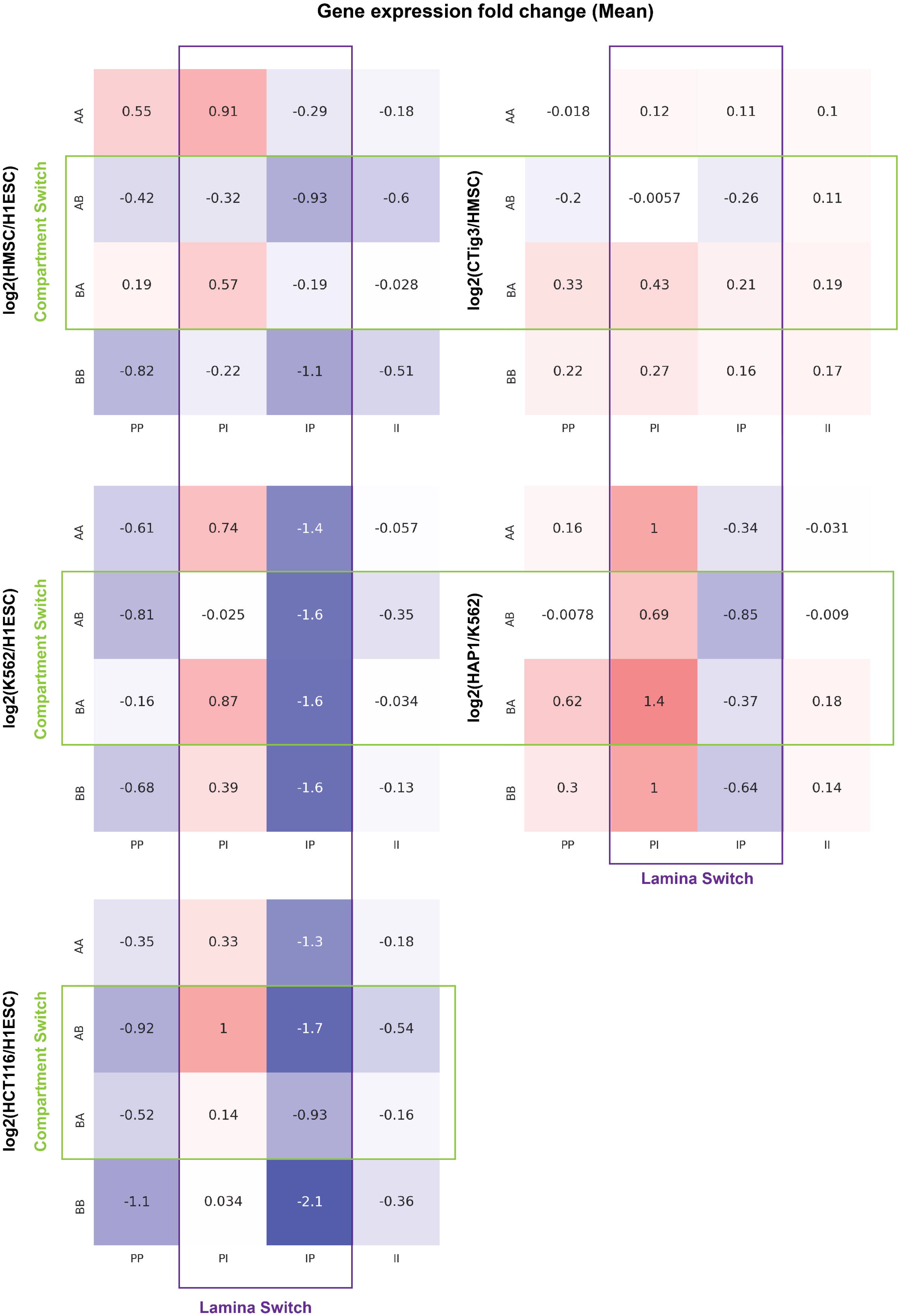
Mean gene expression fold change for different categories of genomic regions based on compartment identity and lamina association. Color shading corresponds to gene expression fold change as indicated by the numbers in each box. Acronyms AB and PI are as defined in Figure 2. Cell type comparisons are indicated to the left of each panel. Green box highlights comparisons where regions switch compartments between cell types while purple box highlights regions that change lamina status.

### The strength of compartment identity correlates with lamina association and vice versa

Through the combined analysis of lamina association and compartmental organization data for different cell types, we identified several megabases of genomic regions which change their lamina association while their compartment identity remains unaltered. Since it is not known how the compartment identity strength (CIS), represented by the magnitude of the compartment analysis eigenvector, is affected by nuclear positioning, we compared the distribution of the CIS values for the regions which show peripheral localization in one cell type and internal in another while keeping their compartment identity unaltered (Fig. 4 and SFig. 4). We found that irrespective of the compartment identity, the CIS values tend to decrease as the regions become more peripheral and increase when regions get internalized (Fig. 4).

**Figure 4.**
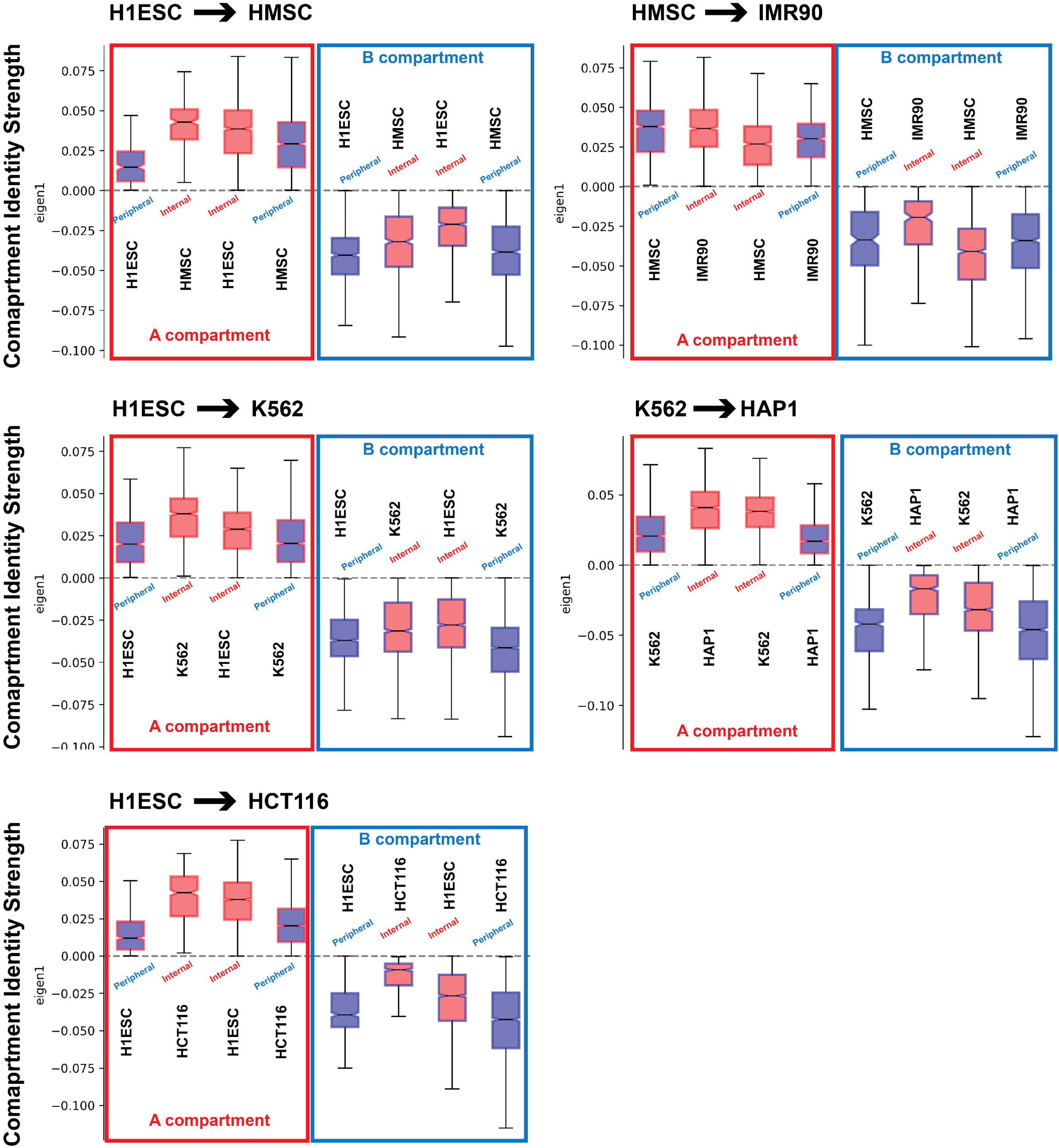
Compartment identity strength shifts with changes in LAD status even when compartment identity remains unchanged. The state of each set of regions is indicated by box plot border color (compartment identity: A = red, B = blue) and shading (LAD = blue, non-LAD = red). Boxplots show minimum, 1^st^ quartile, median, 3^rd^ quartile, and maximum.

In the converse analysis, we considered the strength of the lamin association signal (DamID-seq signal) for regions that change compartment status while their LAD status remains constant between two cell types (SFig. 5). In the majority of cases, the LAD signal becomes stronger when the compartment shifts from A to B. Especially if the region was already classified as lamina associated in the first cell type, a shift to the B compartment often resulted in shifts to stronger lamina association (see H1->HT1080 comparison in SFig 5a). In contrast, in some cell type comparisons, non-LAD regions shifted to even weaker lamina signal despite a change to the B compartment (see H1->Jurkat or K562 comparison in SFig. 5c), suggesting that these B compartments are not mediated by lamin association. For B to A compartment shifts, a similar phenomenon was observed. For regions that were already internal, the compartment shift to A tended to reinforce the lamin status with a shift to even lower lamin association values (see for example comparisons in SFig. 5c). However, LAD-associated regions were less likely to show a decrease in lamin association signal with a shift to the A compartment (see comparisons in SFig. 5a).

### Lamina-associated regions with similar compartment identity exhibit differential histone modifications between cell types

Epigenetic marks are known to be associated with both compartment identity and lamina association^25, 43^45 ^45^. Therefore, we looked into the enrichment of different histone marks in those regions which exhibit similar compartment identity but differential peripheral organization between different cell types. To this end we focused on three different types of broad histone marks: H3K4me1 (active; enhancer), H3K27me3 (facultative heterochromatin), and H3K9me3 (constitutive heterochromatin). Details regarding the calculation of the enrichment of these different histone marks can be found in the **Methods**. We found that genomic regions with similar compartment identity but different lamina association status between cell-types exhibit differential histone modifications in a lineage-specific manner (Fig. 5). For example, in the case of mesenchymal lineage comparisons, we noticed an enrichment of H3K4me1 active mark as the regions move towards the periphery, which holds for both A and B compartment regions. In contrast, for the other cell lineage comparisons, we observed a decrease in the H3K4me1 level as the regions move towards the periphery. Likewise, we noticed a lineage-specific trend for H3K9me3, where a decreased level of this constitutive heterochromatin mark was associated with the regions that move towards the periphery in most of the cell type comparisons except mesenchymal lineage cells, which show the opposite trend. And finally, the H3K27me3 facultative heterochromatin mark most often increases in regions that show a relocation to the periphery while maintaining the same compartment status.

**Figure 5.**
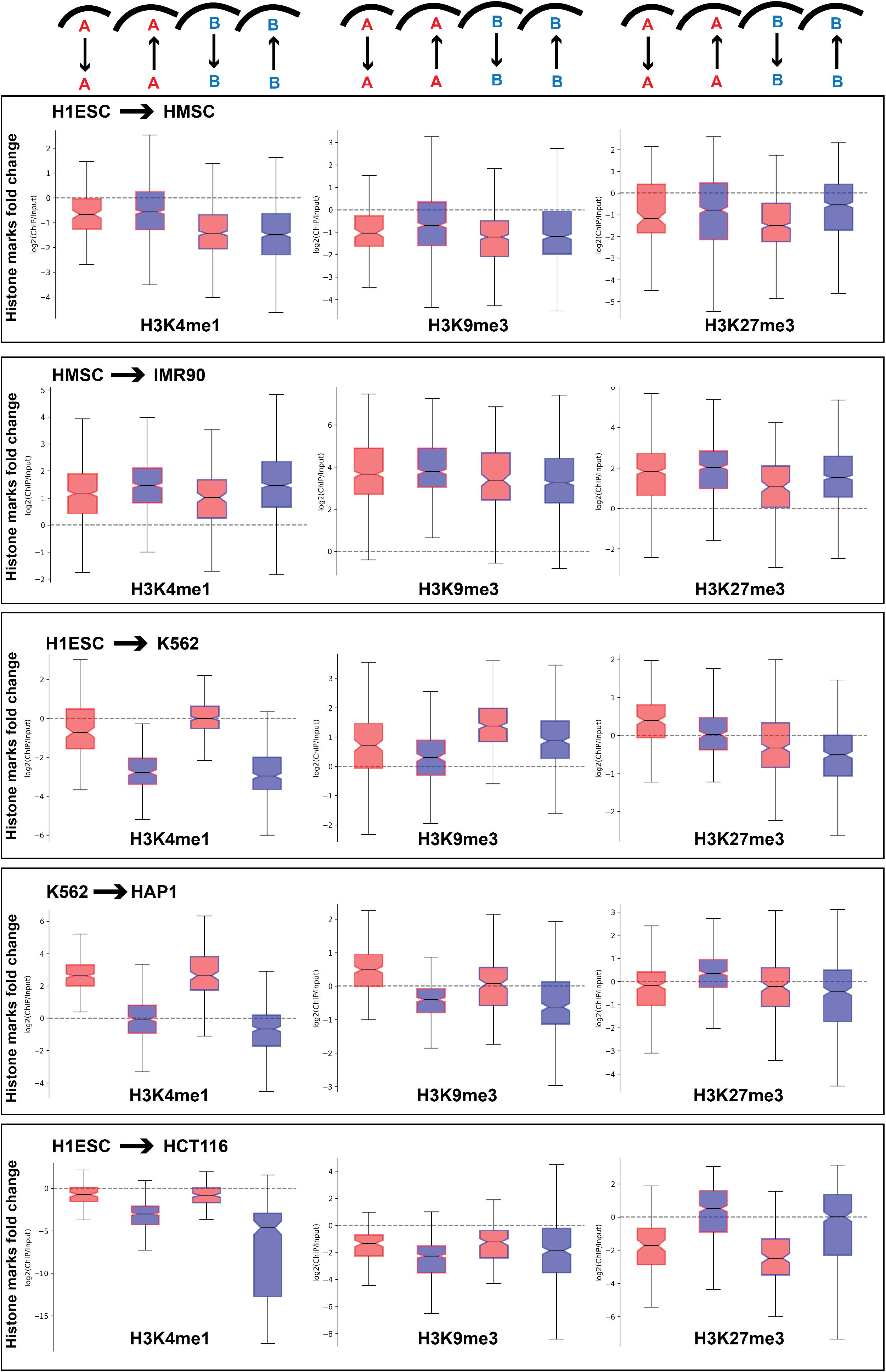
Fold change of different histone marks for the similar compartment identity regions which exhibit differential lamina association between cell types. Shading and labels are as defined in Figure 2. Boxplots show minimum, 1^st^ quartile, median, 3^rd^ quartile, and maximum.

### Lamina association reduces gene expression inducibility after physicochemical treatment

To examine the combined effect of compartmental and spatial localization on gene expression regulation in response to a stimulus, we analyzed transcriptomics data before and after different physicochemical treatments for different cell types with corresponding control lamina association (either DamID or laminB1 ChIP-seq), and compartment identity (Hi-C) data. Using data derived from an acute heat shock exposure of epithelial cells (HCT116), we asked how gene expression regulation was affected by compartment and lamin association in this well-characterized model for rapid and widespread transcriptional response^46^. We found that internal positioning and A compartment status combinatorically increase the likelihood that genes will be upregulated after heat shock (Fig 6a). In contrast, downregulation occurred most frequently in lamina-associated A compartment regions.

**Figure 6.**
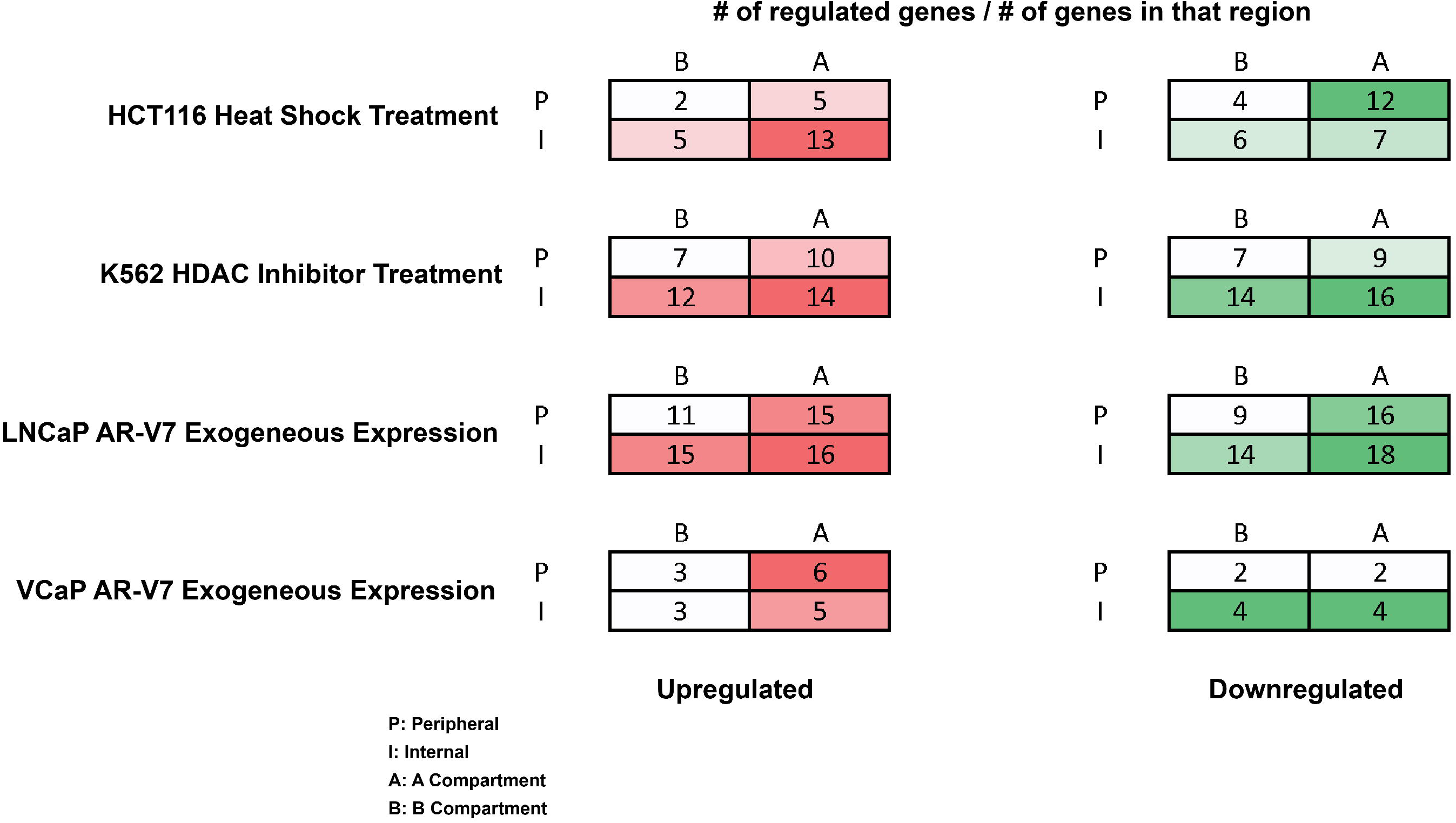
Effect of compartment identity and lamina association on differential gene regulation after different treatments for various cell types. Labels indicate compartment and LAD of each set of regions before treatment and numbers and colors inside the box indicate the percentage of genes in that set of regions that are differentially upregulated (red) or downregulated (green). The differentially regulated genes are selected based on an adjusted p-value cutoff of 0.01.

To evaluate the effect of genome-wide histone deacetylation inhibition, in the context of LAD association and compartmentalization, we analyzed gene expression regulation in K562 cells treated with suberoylanilide hydroxamic acid (SAHA) (Fig. 6b)^47^. This system models the increase in acetylation level of the chromatin at genes regulated by histone deacetylases (HDACs), which can in turn lead to changes in gene expression. In this condition, internal positioning, and A compartment identity combinatorically contributed to differential regulation, for both up- and down-regulation.

Finally, to assess how gene expression regulation can be affected by changes in transcription factor occupancy, we analyzed the effect of exogenous expression of androgen receptor splice variant 7 (AR-V7) in prostate adenocarcinoma LNCaP and bone metastatic VCaP cells^48^. As with heat shock and histone deacetylase inhibitor treatment, both internal positioning and A compartment status increased the amount of gene activation or repression observed in LNCaP cells after AR-V7 induction (Fig. 6c). Gene expression changes were more modest in VCaP cells, with the percentage of upregulated genes correlated with the compartment status. In contrast, the fraction of genes downregulated was more associated with positioning relative to the lamina (Fig. 6d).

## DISCUSSION

The nuclear lamina has a well-recognized function of providing mechanical support to the nucleus and acting as a backbone to genome organization^49–56^. Functionally, chromatin contacts with the nuclear lamina have primarily been regarded as a mechanism for gene expression suppression. By creating a heterochromatic microenvironment underneath the nuclear membrane, the nuclear lamina contributes to silencing of the chromatin regions^6, 57, 58^. Accordingly, genomic regions with heterochromatic properties are often localized to the nuclear periphery and euchromatin regions are found to be primarily located in the nuclear interior. Moreover, following that trend, when a region switches from a euchromatic to a heterochromatic state, it is more likely to assume a more peripheral localization. However, if we set aside the global trends and look systematically along the genome, at significant numbers of genomic regions, switches in either category do not follow the changes in the other. By looking into those different types of incongruent regions, here, we uncovered additional observations about the relationship between genome compartmentalization, lamina association and gene expression. For example, the genomic regions that do not alter their compartment identity between cell types exhibit gene expression changes concordant with their lamina association. Further, this effect is strongest when the lamina association changes in a direction concordant with the pre-existing compartment identity, suggesting that lamina association can reinforce pre-existing chromatin state. In further agreement with this idea, we found a relationship between lamin association strength and compartment identity strength such that, for example, increased lamina association associates with increased compartment identity strength for B compartment regions.

When compartment identity and final lamin association are discordant, some cell types show a strong association between lamin association and gene expression while others do not. This echoes previous observations that radial positioning can be uncoupled from gene expression in some cases^59–62^. Specifically, in the case of the mesenchymal lineage cells, compartment identity is found to be more strongly associated with gene expression regardless of lamin state. For other cell types, the changes in lamina association of those incongruent regions show a stronger association with gene expression. It is possible that in the case of the mesenchymal lineage cells, the genome compartmentalization is relatively decoupled from the lamina association compared to other cell types, as a result of the nuclear geometry in this cell lineage. It can be speculated that due to the flat ellipsoidal geometry of the mesenchymal nuclei, a higher proportion of the genome must be positioned near the periphery regardless of chromatin or compartment state. A nuclear geometry effect may also influence the relationships we observe for the RPE cell type, which behaves more like a mesenchymal cell type in our analyses despite its classification as an epithelial cell. This cell type is often observed to have a small and segmented nucleus, which may again lead to more and different chromosome regions being brought close to the nuclear periphery by geometric constraints.^41, 42^

When considering these results, it is important to keep in mind that the relationships we observe do not prove the directionality of the relationship between these changes. For example, while it could be that a shift of a B compartment region away from the nuclear lamina enables the gene upregulation that we observe in a given cell state transition^63, 64^, it could instead be that the genes in that region were activated first and this led to relocation away from the nuclear lamina, as has been documented in some gene activation experiments^14^. Or, both factors could work in tandem to produce the changes between cell types: the activation of some genes relocates chromatin away from the lamina, which in turn makes the activation of other genes more likely.

Our analyses of gene expression response to physicochemical treatments provide evidence that preexisting lamina association and compartmentalization state influence the ability to alter gene expression in a region. But, while nuclear lamina contacts are often correlated with gene repression, we found that both induction and repression of genes was generally more likely in genomic regions that were located in the A compartment and away from the nuclear lamina. However, specifically in case of heat shock treatment, peripheral A compartment regions exhibit a higher level of downregulation compared to other regions. This may represent regions that are actively transcribed and accessible under normal conditions (thus in the A compartment), but which are poised to be shut down by lamina associated factors in the case of a stimulus.

In our results, while relationships between compartmentalization, lamin association, and gene expression tended to follow principles that often applied across different cell type comparisons, shifts in histone modifications were less consistent between cell types. Some cell lineages showed active and inactive histone modifications changing in concert with lamin association and dissociation while others did not. This may reflect the different relationships that can exist between chromatin modifications and lamin contacts in different regions and contexts. The nuclear lamina can promote alterations in chromatin modifications but also alterations in chromatin state can influence the likelihood of lamina tethering^6^. Since several of our cell lines are cancers derived from the lineages we analyze, it is also important to note that the differences in histone modification relationships we observe may relate to the dysregulation of epigenetic factors common to many cancers.^65–67^ It appears that regardless of divergence of particular combinations of histone marks that are involved in compartment and lamin association status in a given cell type, the resulting spatial organization state produces consistent gene regulatory principles.

## METHODS

### Cell culture

Prostate adenocarcinoma (LNCaP), and bone metastatic (VCaP) cell lines were acquired from the Physical Sciences Oncology Network Bioresource Core Facility, through ATCC (Manassas, VA). These cell lines were cultured according to standard protocols, with subculturing at 80% confluence. LNCaP media was made of RPMI (Gibco;11835030), supplemented to match the ATCC formula (4.5 g/L glucose, 2.383 g/L HEPES, 0.11 g/L sodium pyruvate, 10% FBS). VCaP were cultured in DMEM F12: Ham 1:1 (Gibco; 11-320-033) containing 10% FBS. Media for all cell lines was supplemented with 100 mg/ml penicillin streptomycin (Gibco; 15-140-122).

### Prostate cancer cell line Lamin B1 ChIP-seq

LNCaP and VCaP cells growing under continuous culture conditions in T75 flasks were allowed to reach 90% confluency. LNCaP cells were grown in RPMI (Gibco; 11835030) supplemented with 4.5 g/liter glucose, 2.383 g/liter HEPES, 0.11 g/liter sodium pyruvate and 10% FBS (Corning; 35–010-CV). VCaP cells were cultured in DMEM F12: Ham 1:1 (Gibco; 11–320-033) with 10% FBS. After media removal, the cell monolayer was washed once with HBSS, which was promptly removed by aspiration. Cells were fixed in-situ by incubation in 1% formaldehyde (37% formaldehyde diluted in HBSS) for 10 minutes, under gentle shaking. Crosslinking was quenched by adding glycine to a final concentration of 0.14 M (MP biomedicals ICN19482591), followed by a 5 min incubation at room temperature, with shaking. Flasks were then cooled down on ice for 15 min, after which the formaldehyde solution was aspirated from the plate and substituted with 10 ml of ice-cold HBSS, supplemented with 1X Halt Protease Inhibitor cocktail (Thermo PI78438). Cells were then collected by centrifugation and snap frozen in liquid nitrogen. A replicate flask, cultured under identical conditions, was trypsinized and cells counted, using trypan blue exclusion, to estimate the number of cells per pellet.

Cell pellets were lysed and chromatin was fragmented using a Covaris M220 Sonicator, and the Covaris truChIP Chromatin shearing kit (Covaris; 520154), according to the manufacturers’ instructions, using 1 ml AFA tubes (Covaris). Briefly, cell pellets were thawed on ice and resuspended in 1 ml of 1X lysis buffer, supplemented with protease inhibitors. Samples were incubated on ice for 10 minutes, with gentle vortexing every 2 min. Nuclei were collected by centrifugation at 1700 g for 5 min (at 4 o C). The nuclei pellet was then resuspended in 1 ml of 1X wash buffer (supplemented with protease inhibitors) and collected by centrifugation as before. Samples were then washed twice, using centrifugation as before, with 1 ml of complete shearing buffer. Pellets were resuspended in 1 ml of fresh shearing buffer and transferred into a cold 1 ml AFA tube (Covaris). Sonication conditions were as follows: PIP 75, duty factor 10%, CBP 200, 4 o C, 12 minutes.

To verify sonication, aliquots of time-zero and sonicated samples were treated with RNase A (30 min at 37 o C) and reverse crosslinked overnight with proteinase K via incubation at 65 o C. After reverse crosslinking, DNA was purified using QIAquick PCR Purification Kit (Qiagen; 28104), according to the manufacturer’s instructions. Shearing efficiency was verified via agarose gel electrophoresis.

Chromatin immunoprecipitation was done using the MAGnify Chromatin immunoprecipitation system (Life Technologies, 49-2024), according to the manufacturer’s protocol, diluting the sheared chromatin at a 1:3 ratio in the kit’s dilution buffer and using three reactions per sample. Three micrograms of rabbit polyclonal antibody against Lamin B1 were used, per reaction (Abcam; ab16048). After pulldown and purification, DNA fragments were sonicated for a second time, using the following parameters in a Covaris M220 sonicator: 130 microliter AFA tube, 10 o C, PIP 50%, duty factor 20%, CPB 200, for 45 seconds.

Library preparation and sequencing services were provided by Azenta/Genewiz (South Plainfield, NJ).

### DamID-seq data processing

DamID-seq data were processed as reported in Leemans et al. 2019, except the trimming step^25^. Briefly, at first, bwa mem tool (https://github.com/lh3/bwa) was used to map gDNA reads starting with GATC to a combination of hg19 reference genome with a ribosomal model. For further processing, only the mapped reads having a mapping quality of at least 10 are considered as GATC fragments. Next, the reads were combined into the bins of 40 kb resolution depending on the middle of the GATC fragments and then scaled to 1M reads. For normalization, log2-ratio of the scaled target over the scaled Dam-only bins was calculated with a pseudo count of 1. Furthermore, a Hidden Markov Model was used to call LADs from the normalized values using HMMt R package (https://github.com/gui11aume/HMMt).

### Hi-C data processing

In this study, the HiC-Pro pipeline (https://github.com/nservant/HiC-Pro) was used with default parameters to process all the Hi-C sequencing data. The reads were aligned to hg19 human reference genome^68^. To improve the quality of the contact data, reads from all the replicates of same cell-type or condition were combined together and used for further downstream processing. For all the Hi-C datasets, for each individual chromosome, the contacts were binned at 250 kb resolution and represented in a symmetric square matrix format. Each of the matrices was then further analyzed to obtain the compartmental identity of the chromosome structures at 250 kb resolution using the ‘matrix2compartment.pl’ script from the cworld-dekker (https://github.com/dekkerlab/cworld-dekker) tool suite. Briefly, principal component analysis was performed on the distance-decay normalized contact matrix and A/B compartment identities were assigned based on the sign of the eigenvectors (positive – A and negative – B).

### RNA-seq data processing

For all the RNA-seq data, the fastq reads were first processed for adapter trimming and then for quality trimming using the BBDuk tool (https://github.com/kbaseapps/BBTools)^69^. The parameters used for those adapter and quality trimming steps were “ktrim=r k=23 mink=11 hdist=1” and “qtrim=r trimq=28” respectively. By performing that quality trimming step, reads with quality score lower than 28 were discarded. After having the adapter trimmed and low-quality reads removed, the remaining reads were aligned to human reference genome hg19 with the help of STAR aligner (https://github.com/alexdobin/STAR) and for that step both the ‘--outFilterScoreMinOverLread’ and ‘ – outFilterMatchNminOverLread’ parameters values set to 0.2^70^. Finally, the mapped aligned reads were sorted based on their genomic coordinates and then gene level feature count step was performed on those reads using HTSeq-Counts (https://github.com/simon-anders/htseq)^71^.

### Batch effect removal

The RNA-seq files used in this study were generated at different laboratories using various types of transcriptome profiling techniques. As a result of that the raw gene counts calculated by HTSeq-Counts tool suffer from batch effect, where the samples segregate based on their origin lab and experiment types without showing underlying true biological signal, when analyzed further for downstream processing. To uncover the biological signal, the ComBat-seq tool (https://github.com/zhangyuqing/ComBat-seq) was applied on the raw counts for the batch effect adjustments^72^. Un this step, the files originated from the same laboratory were assigned same batch number.

### Differential expression analysis

Differential gene expression analysis was performed between different condition samples of the same cell type (for example, before and after AR7 induction in LNCaP and VCaP cells) using DESeq2 (https://bioconductor.org/packages/release/bioc/html/DESeq2.html)^73^. To identify significantly differentially regulated genes, we use an adjusted p-value cutoff of 0.01.

### Histone ChIP-seq data processing

BBDuk tool (https://github.com/kbaseapps/BBTools) was used to process all the histone ChIP-seq data for adapter removal and quality trimming^69^. After performing those steps, the reads were mapped to hg19 human reference genome using STAR aligner (https://github.com/alexdobin/STAR)^70^. Finally, the peaks were called from the aligned data using MACS2 (https://pypi.org/project/MACS2/) peak calling software^74^. Since, in this study, we are mainly focusing on broad histone marks, here we used MACS2 broad peak calling parameters “-p 1e-2 --broad --nomodel --shift 0 --keep-dup all -B --SPMR”. In addition to all those parameters, we also supplied the estimated fragment length obtained from the ChIP-seq data as “--extsize” parameter. To calculate the estimated fragment length, ‘run_spp.R’ script from the phandompeakqualtools (https://github.com/kundajelab/phantompeakqualtools) was used^75, 76^. Once we had the broad peak locations for a particular histone mark, the enrichment of that mark for different genomic regions was measured by calculating the base pair overlaps using the ‘intersect’ function of bedtools (https://bedtools.readthedocs.io/en/latest/index.html)^77^.

### LaminB1 ChIP-seq data processing

LaminB1 ChIP-seq files were processed first using the BBDuk tool (https://github.com/kbaseapps/BBTools) for adapter removal and quality trimming^69^. After that, STAR aligner (https://github.com/alexdobin/STAR)^70^ was used to map the reads to the hg19 genome. Once, both the target and input mapped reads were mapped, they were binned at 40 kb resolution and the log2-ratio of the target over the input bins was calculated using ‘bamCompare’ function of deepTools (https://deeptools.readthedocs.io/) with parameters “--operation log2 -bs 40000 --ignoreDuplicates --minMappingQuality 30 --scaleFactorsMethod SES --effectiveGenomeSize 2864785220”^78^. The signal is then further smoothed using a rolling window averaging (120 kb) followed by LAD prediction by hidden markov model as described in the DamID-seq data processing.

### Hierarchical clustering of genome-wide lamina association and compartmental organization data

For both types of data, we performed hierarchical clustering to cluster the cell types based on similarity of the respective data type. To do that, we first calculated the pairwise similarity between all the 11 cell types using Pearson’s correlation. We further converted the similarity data into distance data by subtracting the values from 1. Following that, finally we used the distance data for hierarchical clustering using ‘ward’ linkage.

### Combine lamina association and compartmental organization data

We combined the LAD and compartment identity data together for each cell type. For that, we used the ‘intersect’ function of bedtools (https://bedtools.readthedocs.io/en/latest/index.html) to combine genomewide LAD and compartment identity data together with a minimum 50% overlap as a fraction of LAD data^77^. This allowed us to categorize the genomic regions based on whether they belong to LAD or not and also simultaneously whether those regions are part of A or B compartment.

## ACKNOWLEDGEMENTS

The authors would like to thank all the members of McCord lab for their fruitful suggestions in developing the research project.

## DATA AVAILABILITY

Datasets used for this project were obtained from NCBI GEO, EMBL ENA, 4DN Data Portal and ENCODE repositories and their accession IDs are given in Supplementary Table 1. Newly generated LaminB1 ChIP data for VCaP and LNCaP are available at GEO with accession number GSE213032.

## AUTHOR CONTRIBUTIONS

P.D. conceived the study and performed data analysis and prepared figures. R.S.M. performed the Lamin B1 ChIP experiments and assigned cell types to lineages. R.P.M. supervised the project, provided funding, and prepared figures. P.D. and R.P.M. wrote the manuscript with input from R.S.M.

## SUPPLEMENTARY MATERIAL

The supplementary material associated with the paper can be found and accessed online.

## SUPPLEMENTARY TABLE AND FIGURES LEGENDS

**Supplementary Table 1**. NCBI GEO, EMBL ENA, 4DN Data Portal and ENCODE accession IDs for the Hi-C, DamID-seq, RNA-seq and ChIP-seq data from thirteen different cell-types.

**Supplementary Figure 1**. **Proportion of LAD and compartment type for each cell type**. a) Percentages of peripheral and internal regions based on lamina association for different cell types. b) Percentages of A and B compartment regions based on genome organization data for different cell types.

**Supplementary Figure 2**. **Gene expression changes with alteration in lamina association for similar compartment identity genomic regions**. Labels, coloring, and organization are as described in Figure 2, for different cell type comparisons.

**Supplementary Figure 3**. **Mean gene expression fold change between cell types for different categories of genomic regions based on compartment identity and lamina association.** Labels, coloring, and organization are as described in Figure 3, for different cell type comparisons.

**Supplementary Figure 4**. **Compartment identity strength shifts with changes in LAD status even when compartment identity remains unchanged.** Labels, coloring, and organization are as described in Figure 4, for different cell type comparisons.

**Supplementary Figure 5**. **Lamina association strength shifts with changes in compartment status.** Cell type comparisons shown along the a) epithelial lineage, b) mesenchymal lineage, and c) hematopoietic lineage. Labels, coloring, and organization are as described in Figure 4.

**Supplementary Figure 6**. Fold change of different histone marks for the similar compartment identity regions which exhibit differential lamina association between cell types. Labels, coloring, and organization are as described in Figure 5, for different cell type comparisons.

