## Supplementary Figures and Table for "Differential contributions of nuclear lamina association and genome compartmentalization to gene regulation"

**Supplementary Table 1**. NCBI GEO, EMBL ENA, 4DN Data Portal and ENCODE accession IDs for the Hi-C, DamID-seq, RNA-seq and ChIP-seq data from thirteen different cell-types. RC = replicates combined for analysis, R1 = replicate 1 used for analysis. Specific conditions chosen from each accession number indicated where needed in parentheses.

| Cell type | Hi-C | DamID-seq | RNA-seq | H3K4me1 ChIP-seq | H3K27me3 ChIP-seq | H3K9me3 ChIP-seq |
|---|---|---|---|---|---|---|
| H1ESC | 4DNESRJ8KV4Q (RC) | 4DNESOFQR5FS (LaminB1; RC) | ENCSR588EJX (RC) | ENCSR991BTY (R1) | ENCSR687FDK (R1) | ENCSR395USV (R1) |
| HMSC | GSE140782 (RC) | GSE146387 (Emerin; RC) | GSE108638 (RC) | ENCSR065HOR (R1) | ENCSR832JVP (R1) | ENCSR746CUY (R1) |
| IMR90 | X | X | X | GSE106146 (Proliferating; R1) | GSE135088 (Growing; R1) | GSE53332 (OIS_Proliferating; R1) |
| WI38 | PRJEB8073 (Cycling; RC) | X | X | X | X | X |
| Tig3 | X | GSE76594 (LaminB1; Cycling; RC) | GSE75643 (RC) | X | X | X |
| HFFc6 | GSE165894 (RC) | 4DNESXZ4FW4T (LaminB1; RC) | 4DNESFH3EHTU (RC) | X | X | X |
| U2OS | GSE137469 (RC) | 4DNESB2HMY7D (LaminB1; RC) | GSE131571 (RC) | GSE73742 (R1) | GSE35573 (R1) | GSE69809 (R1) |
| HCT116 | GSE133023 (RC) | 4DNES24XA7U8 (LaminB1; RC) | GSE162160 (RC) | ENCSR161MXP (R1) | ENCSR810BDB (R1) | ENCSR179BUC (R1) |
| HT1080 | GSE117582 (RC) | GSE87148 (LaminB1; RC) | ENCSR535VTR (RC) | X | GSE153869 (R1) | GSE135580 (R1) |
| RPE | ENCSR846DUS (RC) | 4DNESHGTQ73M (LaminB1; RC) | GSE163315 (RC) | GSE163315 (R1) | X | GSE163315 (R1) |
| K562 | ENCSR545YBD (RC) | 4DNES6SM8UGO (LaminB1; RC) | ENCSR615EEK (RC) | ENCSR000AKS (R1) | ENCSR000AKQ (R1) | ENCSR000APE (R1) |
| HAP1 | ENCSR390HMC (RC) | 4DNESUK5H9Y8 (LaminB1; RC) | GSE160383 (RC) | ENCSR450JTP (R1) | ENCSR891YFI (R1) | ENCSR460TQV (R1) |
| Jurkat | GSE122958 (RC) | GSE94971 (LaminB1; Resting; RC) | GSE130140 (RC) | GSE119439 (R1) | GSE100693 (R1) | GSE162605 (R1) |

**Supplementary Figure 1. Proportion of LAD and compartment type for each cell type**. a) Percentages of peripheral and internal regions based on lamina association for different cell types. b) Percentages of A and B compartment regions based on genome organization data for different cell types.

### Supplementary Figure 1.

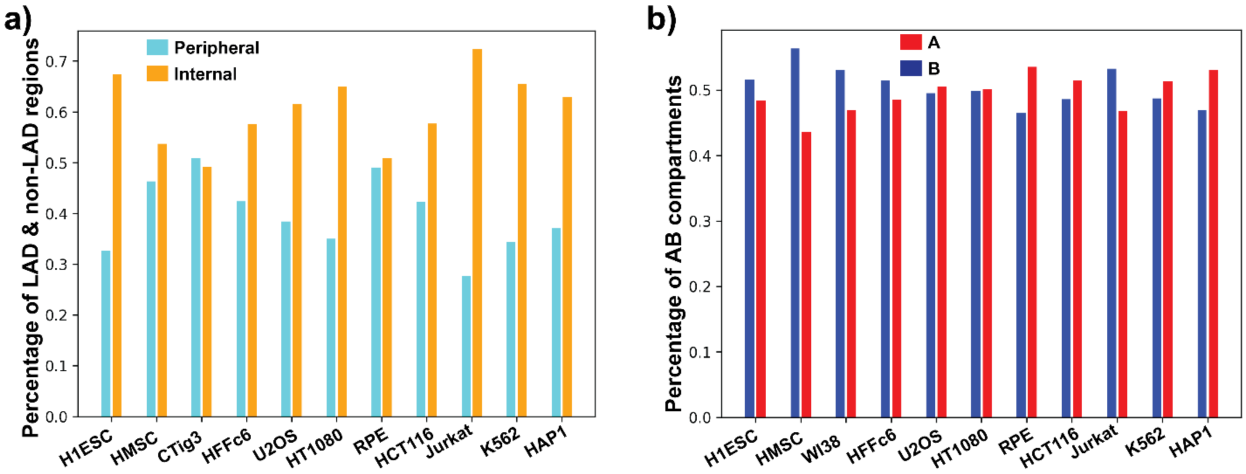

**Supplementary Figure 2. Gene expression changes with alteration in lamina association for similar compartment identity genomic regions**. Labels, coloring, and organization are as described in Figure 2, for different cell type comparisons.

#### Supplementary Figure 2.

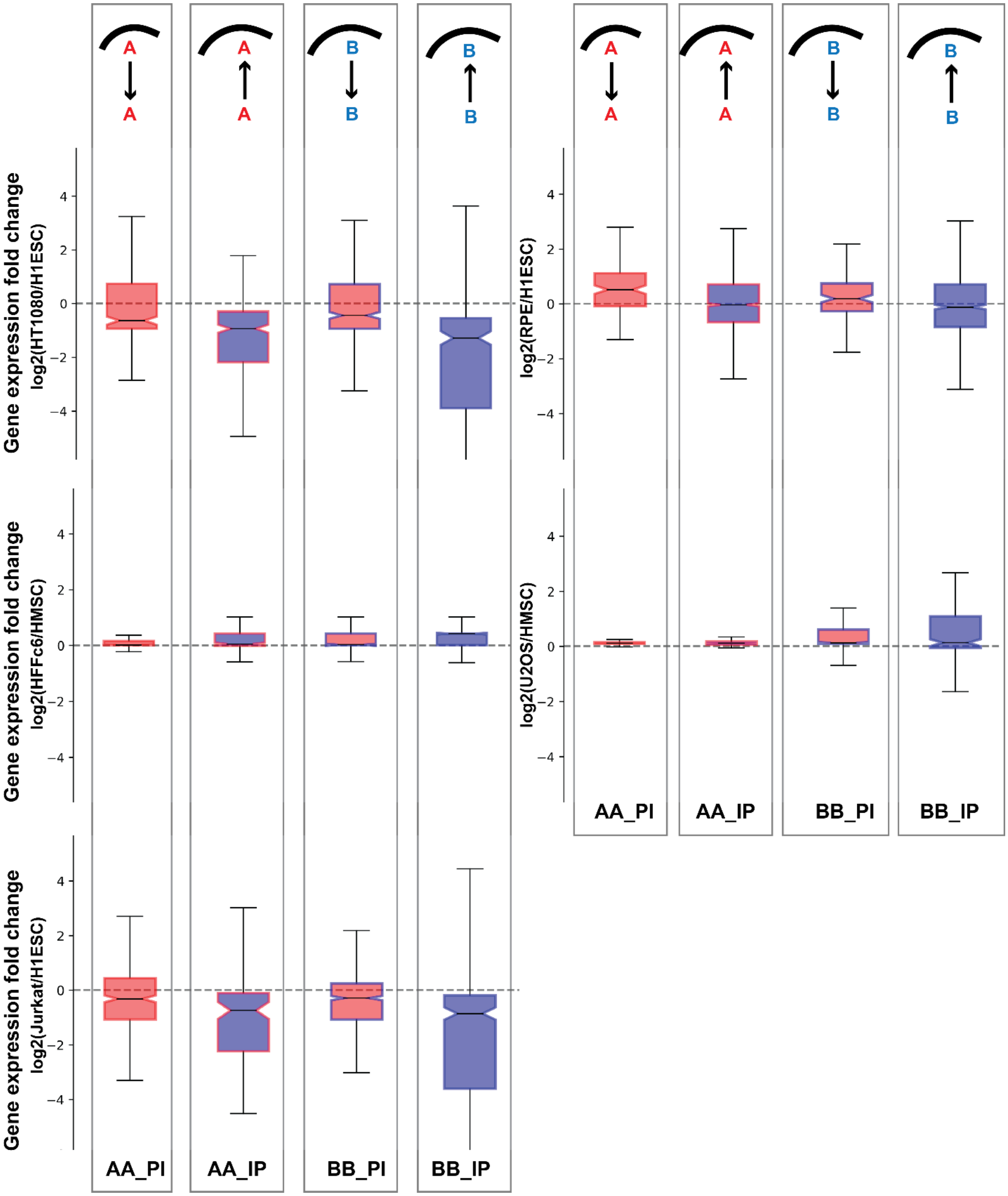

**Supplementary Figure 3. Mean gene expression fold change between cell types for different categories of genomic regions based on compartment identity and lamina association.** Labels, coloring, and organization are as described in Figure 3, for different cell type comparisons.

### Supplementary Figure 3.

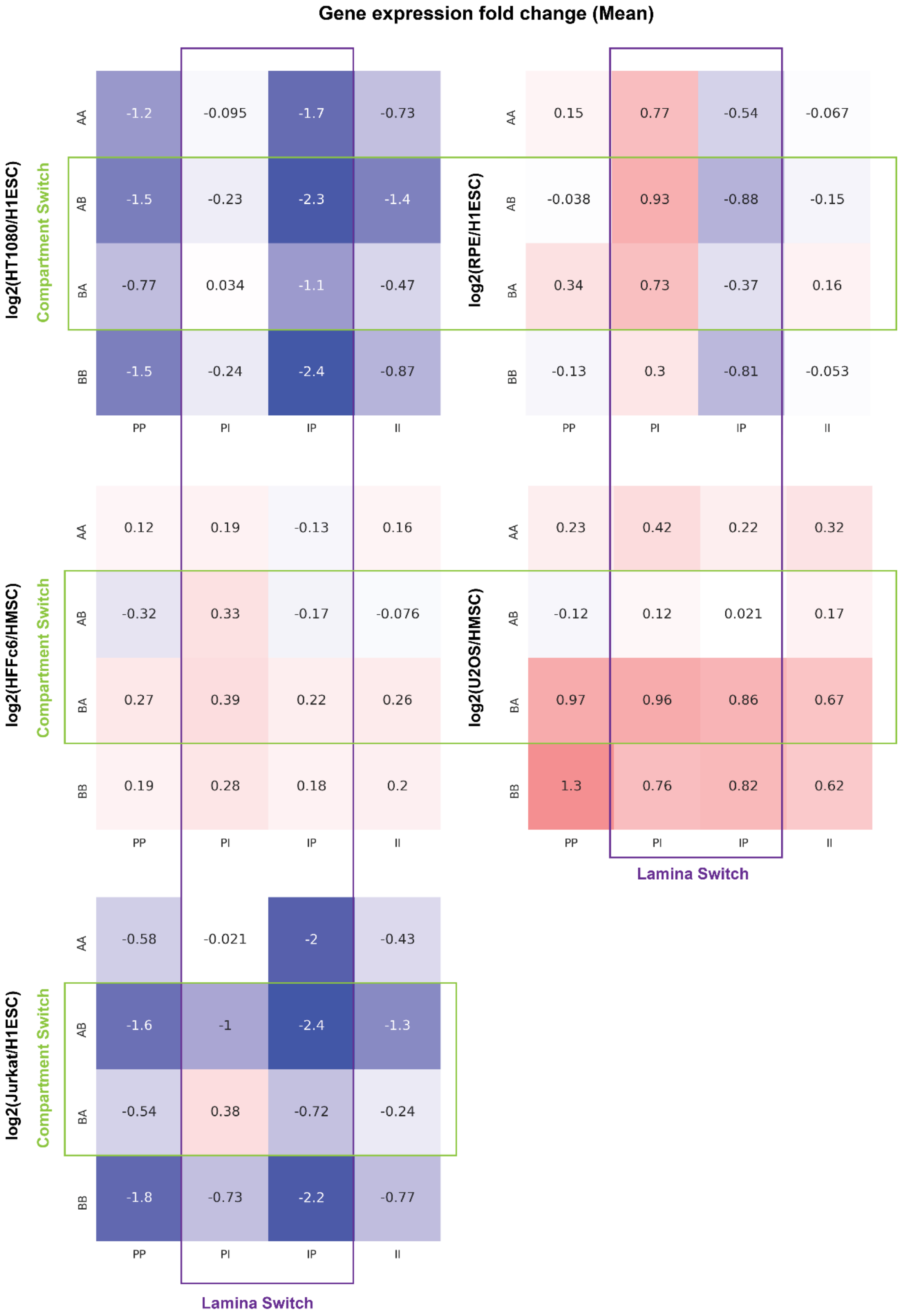

**Supplementary Figure 4. Compartment identity strength shifts with changes in LAD status even when compartment identity remains unchanged.** Labels, coloring, and organization are as described in Figure 4, for different cell type comparisons.

### Supplementary Figure 4.

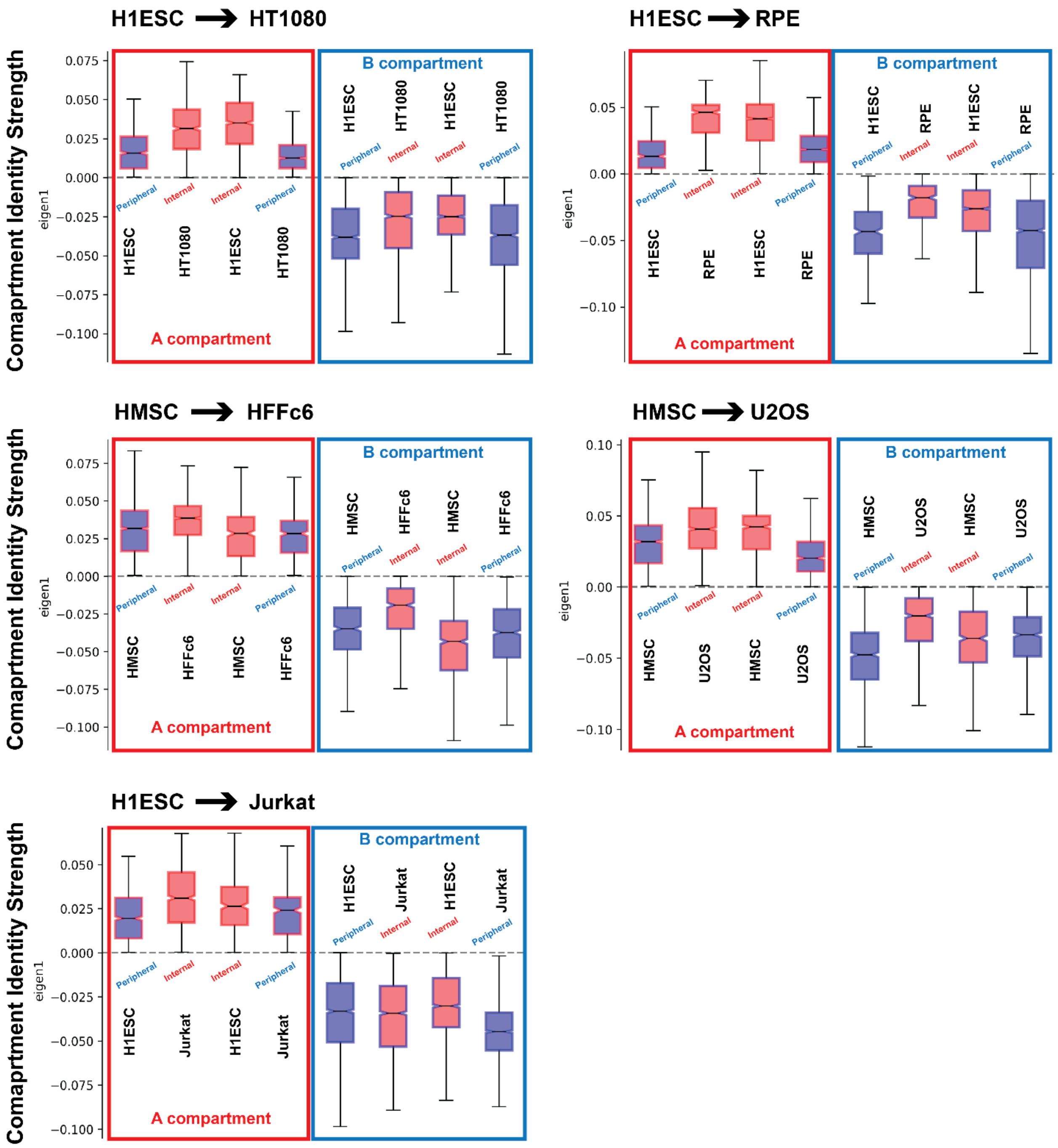

**Supplementary Figure 5. Lamina association strength shifts with changes in compartment status.** Cell type comparisons shown along the a) epithelial lineage, b) mesenchymal lineage, and c) hematopoietic lineage. Labels, coloring, and organization are as described in Figure 4.

### Supplementary Figure 5a.

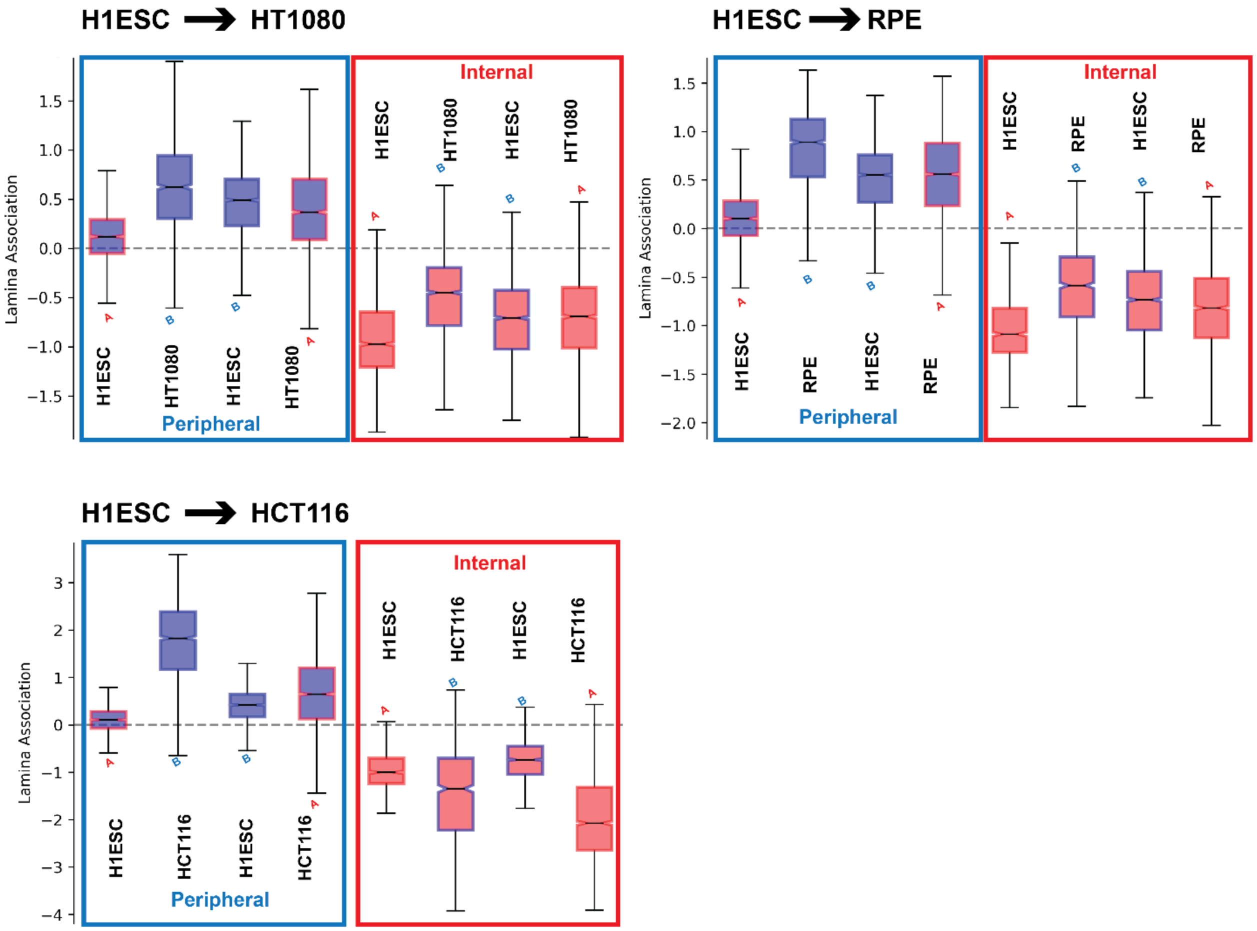


**Lamina Association = log2(Dam Protein / Dam only)**

#### Supplementary Figure 5b.

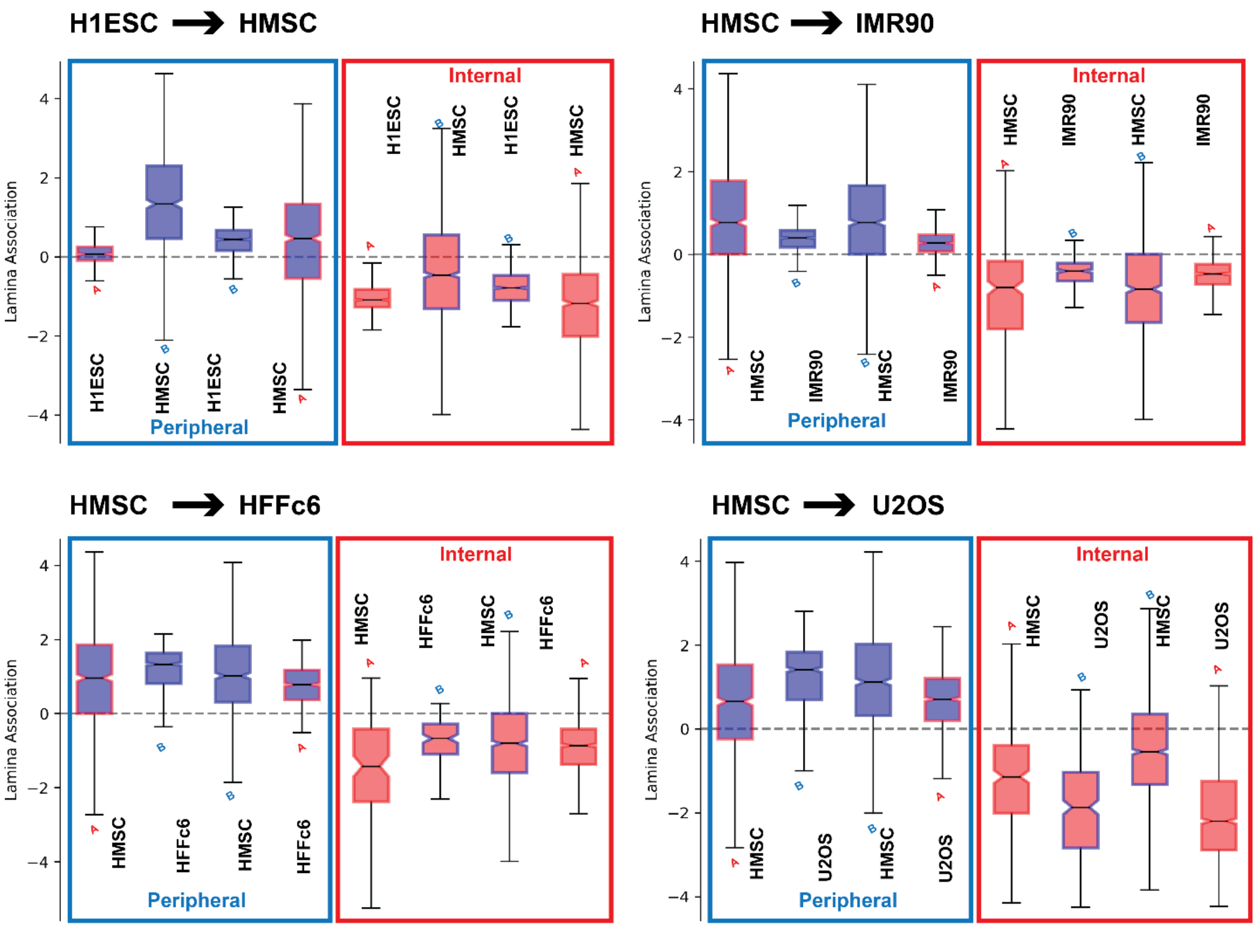


Lamina Association = log2(Dam Protein / Dam only)

### Supplementary Figure 5c.

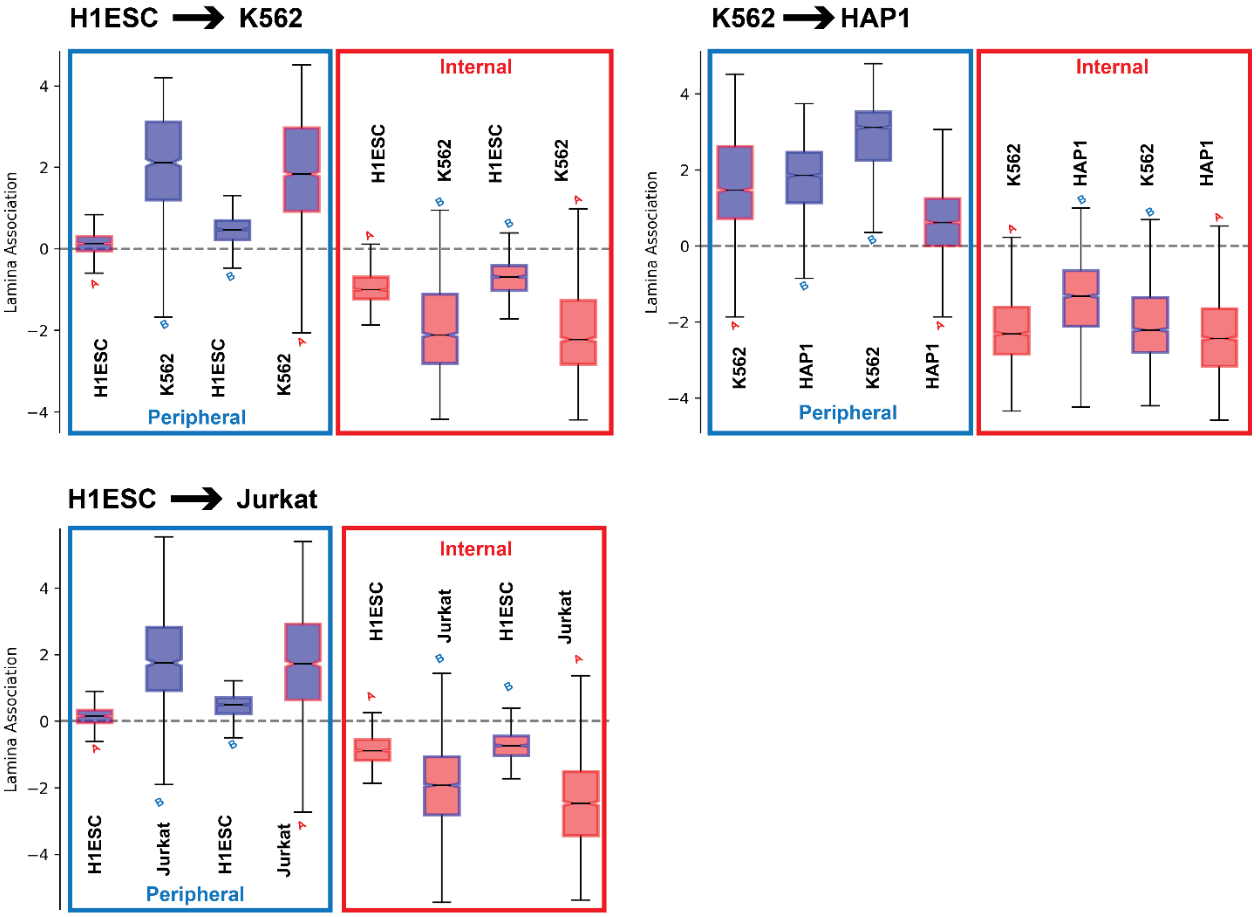


Lamina Association = log2(Dam Protein / Dam only)

**Supplementary Figure 6**. Fold change of different histone marks for the similar compartment identity regions which exhibit differential lamina association between cell types. Labels, coloring, and organization are as described in Figure 5, for different cell type comparisons.

#### Supplementary Figure 6.

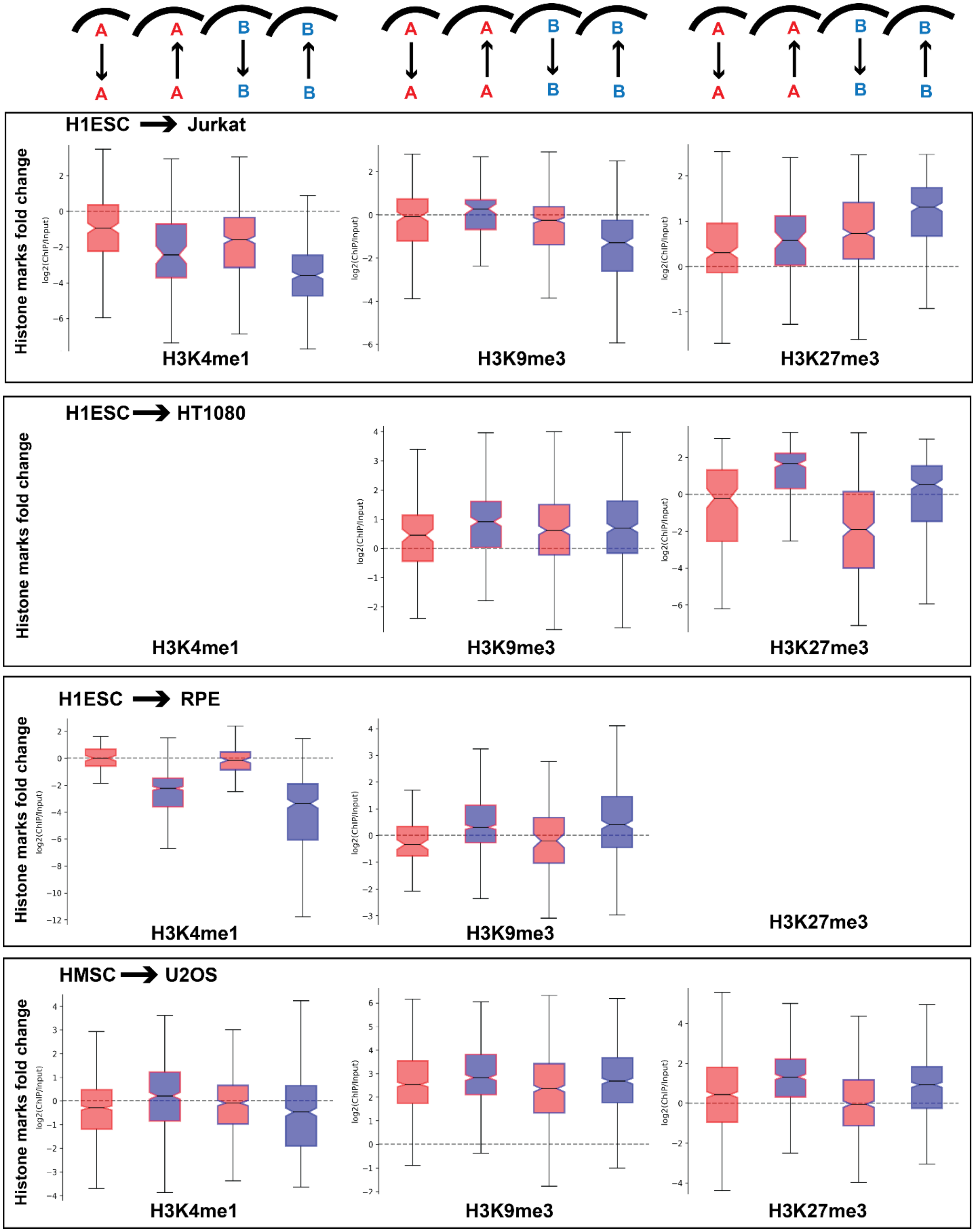
